# The Avian RNAseq Consortium: a community effort to annotate the chicken genome

**DOI:** 10.1101/012559

**Authors:** Jacqueline Smith, David W. Burt, the Avian RNAseq Consortium

## Abstract

Here we describe how members of the chicken research community have come together as the “Avian RNAseq Consortium” to provide chicken RNAseq data with a view to improving the annotation of the chicken genome. The data was used by Ensembl in their release 71 gene_build to help provide the most up-to-date annotation of Galgal4, which is still in current use. The data has also been used to predict many ncRNAs, particularly lncRNAs, and it continues to be used as a resource for annotation of other avian genomes along with continued gene discovery in the chicken. This article is submitted as part of the Third Report on Chicken Genes and Chromosomes which will be published in Cytogenetic and Genome Research

---

Publication of the chicken genome sequence in 2004 (International Chicken Genome Sequencing Consortium 2004) highlighted the beginning of a revolution in avian genomics. Progression of DNA sequencing technologies and data handling capabilities has also meant that genome sequencing and assembly is now a relatively simple, fast and inexpensive procedure. The success seen with the chicken genome was soon followed by the completion of the zebra finch genome (Warren et al., 2010), an important model for neurobiology (Clayton et al., 2009), again based on Sanger sequencing. In recent years the rapid advances in Next Generation Sequencing (NGS) technologies, hardware and software have meant that many more genomes can now be sequenced faster and cheaper than ever before (Metzker, 2010). The first avian genome to be sequenced by NGS methods was the turkey (Dalloul et al., 2010), which was also integrated with genetic and physical maps thus providing an assembly of high quality, even at the chromosome level. Recently, NGS has been used to sequence the genomes of a further 42 avian species, as part of the G10K initiative (Genome 10K Community of Scientists, 2009). In addition there have also been 15 other genome assemblies recently published, each with a focus on a unique aspect of avian biology, including the Japanese Quail (domestication; Kawahara-Miki et al., 2013), Puerto Rican parrot (speciation; Oleksyk et al., 2012), Scarlet Macaw (speech, intelligence and longevity; Seabury et al., 2013), Medium and Large Ground Finches (speciation; Parker et al., 2012; Rands et al., 2013), Collared and Pied flycatchers (speciation; Ellegren et al., 2012), Peregrine and Saker Falcons (predatory lifestyle; Zhan et al., 2013), rock pigeon (domestication; Shapiro et al., 2013), the Ground tit (adaptation to high altitude; Cai et al., 2013) and the Northern Bobwhite (population history; Halley et al., 2014). Through November 2014 there are currently 57 avian genome sequences completed, either published or in press (Table 1). A new project, B10K (web.bioinfodata.org/B10K), proposes sequencing all avian genomes; this would include all 40 orders, 231 families, 2,268 genera and 10,476 species of birds. The chicken genome remains the best described genome and is used as a reference upon which the annotations of other assemblies are based. Assembly and annotation of the genome continues to improve. However, gaps and unaligned regions remain (particularly for some of the smallest micro-chromosomes), which can cause practical problems in the analysis and annotation of important loci, especially for those representing gene families. Other approaches, such as long reads generated by Pacific Biosciences (PacBio) sequencing, chromosome sorting and optical maps are being used to resolve these assembly issues (Warren and Burt, personal communications). Specific genome features also require further study; for example, non-coding RNAs, annotation of rare transcripts, confirmation of alternatively spliced transcripts, mapping of transcription start sites and identification of conserved regions. One method by which some of these goals can be achieved is through analysis of transcriptomic sequence data, or ‘RNAseq’ data.

**Table 1:** Avian species with sequenced genomes

| Table 1: Avian species with sequenced genomes |  |  |  |  |  |  |
| --- | --- | --- | --- | --- | --- | --- |
| BIRD_Abbreviation | BIRD_Latin_Name | BIRD_Common_Name |  | BIRD_Abbreviation | BIRD_Latin_Name | BIRD_Common_Name |
| ACACH | Acanthisitta chloris | Rifleman |  | GALGA | Gallus gallus | Chicken |
| AMAVI | Amazona vittata | Puerto Rican parrot |  | GAVST | Gavia stellata | Red-throated loon |
| ANAPL | Anas platyrhynchos domestica | Pekin duck |  | GEOFO | Geospiza fortis | Medium groundfinch |
| ANOCA | Anolis carolinensis | Carolina anole |  | GEOMA | Geospiza magnirostris | Large ground finch |
| APAVI | Apaloderma vittatum | Bar-tailed trogon |  | HALAL | Haliaeetus albicilla | White-tailed eagle |
| APTFO | Aptenodytes forsteri | Emperor penguin |  | LEPDI | Leptosomus discolor | Cuckoo roller |
| ARAMA | Ara macao | Scarlet macaw |  | MANVI | Manacus vitellinus | Golden-collared manakin |
| BALRE | Balearica regulorum gibbericeps | Grey crowned crane |  | MELGA | Meleagris gallopavo | Wild turkey |
| BUCRH | Buceros rhinoceros silvestris | Rhinoceros hornbill |  | MELUN | Melopsittacus undulatus | Budgerigar |
| CALAN | Calypte anna | Anna's hummingbird |  | MERNU | Merops nubicus | Northern Carmine bee-eater |
| CAPCA | Caprimugus Carolinensis | Chuck-will's widow |  | MESUN | Mesitornis unicolor | Brown mesite |
| CARCR | Cariama cristata | Red-legged seriema |  | NESNO | Nestor notabilis | Kea |
| CATAU | Cathartes aura | Turkey vulture |  | NIPNI | Nipponia nippon | Crested ibis |
| CHAPE | Chaetura pelagica | Chimney swift |  | OPHHO | Ophisthocomus hoazin | Hoatzin |
| CHAVO | Charadrius vociferus | Killdeer |  | PELCR | Pelecanus crispus | Dalmatian pelican |
| CHLUN | Chlamydotis undulata | Houbara bustard |  | PHACA | Phalacrocorax carbo | Great cormorant |
| COLLI | Columba livia | Rock pigeon |  | PHALE | Phaethon lepturus | White-tailed tropicbird |
| COLST | Colius striatus | Speckled mousebird |  | PHORU | Phoenicopterus ruber | American flamingo |
| COLVI | Colinus virginianus | Northern Bobwhite |  | PICPU | Picoides pubescens | Downy woodpecker |
| CORBR | Corvus brachyrhynchos | American crow |  | PODCR | Podiceps cristatus | Great crested grebe |
| COTJA | Coturnix japonica | Japanese quail |  | PSEHU | Pseudopodoces humilis | Ground tit |
| CUCCA | Cuculus canorus | Common cuckoo |  | PTEGU | Pterocles gutturalis | Yellow-throated sandgrouse |
| EGRGA | Egretta garzetta | Little egret |  | PYGAD | Pygoscelis adeliae | Adelie penguin |
| EURHE | Eurypyga helias | Sunbittern |  | STRCA | Struthio camelus | Ostrich |
| FALCH | Falco cherrug | Saker falcon |  | TAEGU | Taeniopygia guttata | Zebra finch |
| FALPE | Falco peregrinus | Peregrine falcon |  | TAUER | Tauraco erythrolophus | Red-crested turaco |
| FICAL | Ficedula albicollis | Collared flycatcher |  | TINMA | Tinamus major | Great tinamou |
| FICHY | Ficedula hypoleuca | Pied flycatcher |  | TYTAL | Tyto alba | Barn owl |
| FULGL | Fulmarus glacialis | Northern fulmar |  |  |  |  |

With a view to addressing some of these issues, we decided to collect as much RNAseq data from the chicken research community as possible. This was the beginning of what we have termed ‘The Avian RNAseq Consortium’. Since the start of the Consortium at the end of 2011, it now includes 48 people from 27 different institutions (Figure 1) who have contributed to the effort to create a detailed annotation of the chicken genome by either providing RNAseq data or by helping to analyse the combined data.

**Figure 1:**
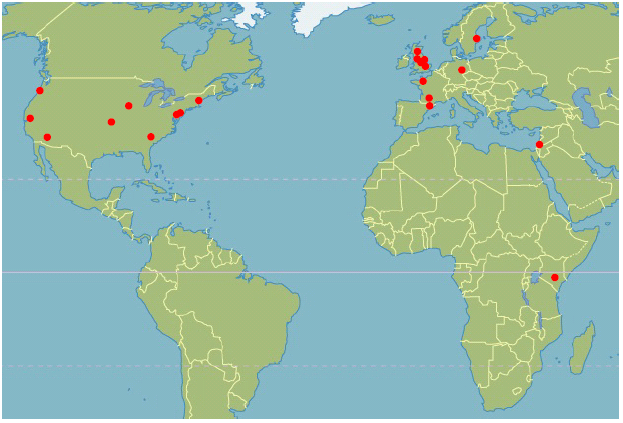
Worldwide locations of current RNAseq consortium members.

We currently have 21 different data sets (representing more than 1.5 Tb of data) with more data being added (Table 2 and Figure 2). These data represent transcriptome sequences from many different chicken tissues and from many different experimental conditions, including several infection/disease cases. These data were submitted to public archives, collected at The Roslin Institute and then passed on to the Ensembl team who used the information to help annotate the latest chicken genome assembly, Galgal4 as part of Ensembl release 71 (April 2013) (Table 3). This new annotation includes 15,495 protein coding genes, 1,049 micro RNAs, 456 non-codingRNAs and 42 pseudo genes. This gene_build is primarily concerned with coding genes, but there are many more non-coding genes which remain unannotated. Consortium members have analysed the RNAseq data for long non-coding RNAs (lncRNAs) [*manuscript in preparation*], snoRNAs (Gardner et al., 2014) and other features of interest. Around 14,000 potential long non-coding RNA genes have thus far been identified from the RNAseq data. Ensembl release 71 marked a significant update in the annotation of the chicken genome with gene models based on experimental data. Table 4 shows how this gene_build was the first to use the Galgal4 assembly and, through the use of RNAseq data, was able to help remove assembly errors and reduce the number of predicted gene transcripts by identifying incorrectly predicted genes from previous builds and improving identification of short ncRNAs. The significance of this community effort is indicated by the fact that the current Ensembl 77 gene set has not changed since Ensembl release 71, with only difference being reflected in the total number of base pairs. This is due to the correction of one particular scaffold on the Z chromosome (which was reflected in Ensembl release 74).

**Table 2:** Details of RNAseq data sets

| Table 2: Details of RNAseq data sets |  |  |  |
| --- | --- | --- | --- |
| Data set | Description of data | Reads (bp) | Sequencing* |
| 1. Antin | Whole embryo | 35 | Illumina SE |
| 2. Blackshear | LPS stimulated macrophages v control CEFs | 51 | Illumina PE |
| 3. Burgess/McCarthy | miRNA from various RJF tissues (adrenal gland, adipose, cerebellum, cerebrum, testis, ovary, heart, hypothalamus, kidney, liver, lung, breast muscle, sciatic nerve, proventriculus, spleen) | 50 | Illumina SE |
| 4. Burt/Smith | Spleen: Infectious Bursal Disease Virus infected v control | 36 | Illumina SE |
| 5. | Lung and ileum: Avian influenza infected v control (high path H5N1 and low | 36 | Illumina SE |
| 6. | Lung short read data | 25 | Illumina SE |
| 7. de Koning/Dunn/McCormack | Bone from 70wk old Leghorns | 100 | Illumina PE |
| 8. Frésard/Pitel | Brain from epileptic v. non-epileptic birds | 380-400 | Roche 454 |
| 9. | Pooled whole embryos (stage HH26) | 100 | Illumina PE |
| 10. Froman/Rhoads | Testes: roosters with high mobility sperm v low mobility sperm | 35 | Illumina SE |
| 11. Garceau/Hume | Embryo, DF1 cell line and bone marrow derived macrophages | 100 | Illumina PE |
| 12. Hanotte/Kemp/Noyes/Ommeh | Newcastle Disease Virus infection v control (trachea and lung epithelial cells) | 50 | SOLiD SE |
| 13. Häslér/Oler/Muljo/Neuberger | DT40 cells | 60 | Illumina PE |
| 14. Kaiser | Bone marrow derived dendritic cells from 6 weeks old birds (Control, DCs +LPS) Bone marrow derived macrophages from 6 weeks old birds (Control, BMDMs +LPS) Heterophils isolated from blood of day-old chicks (Control, DCs +LPS) | 100 | Illumina PE |
| 15. Lagarrigue/Roux | Abdominal adipose tissue and liver tissue from 14wk old broilers | 100 | Illumina PE |
| 16. Lamont | Livers of eight individual, 28-day-old broiler males - 4 control; 4 heat-stressed | 100 | Illumina SE |
| 17. Munsterberg/Pais | Somites injected with anti-mir206 v. non-injected | 50 | Illumina PE |
| 18. Schmidt | Tissues from heat stressed and control birds (liver, brain, spleen, thymus, bursa, kidney, ileum, jejunum, duodenum, ovary, heart, breast, monocyte | 42-50 | Illumina SE |
| 19. Schwartz/Ulitsky | Whole embryo stages - HH4/5; HH11; HH14/15; HH21/22; HH25/26; HH32; HH36 - Stranded | 80/100 | Illumina PE |
| 20. Skinner | Chicken embryo fibroblasts | 100 | Illumina PE |
| 21. Wang/Zhou | Lung from Fayoumi and Leghorn birds - control and H5N3 infected | 75 | Illumina SE |
| * - SE: single end; PE: paired end |  |  |  |

**Figure 2:**
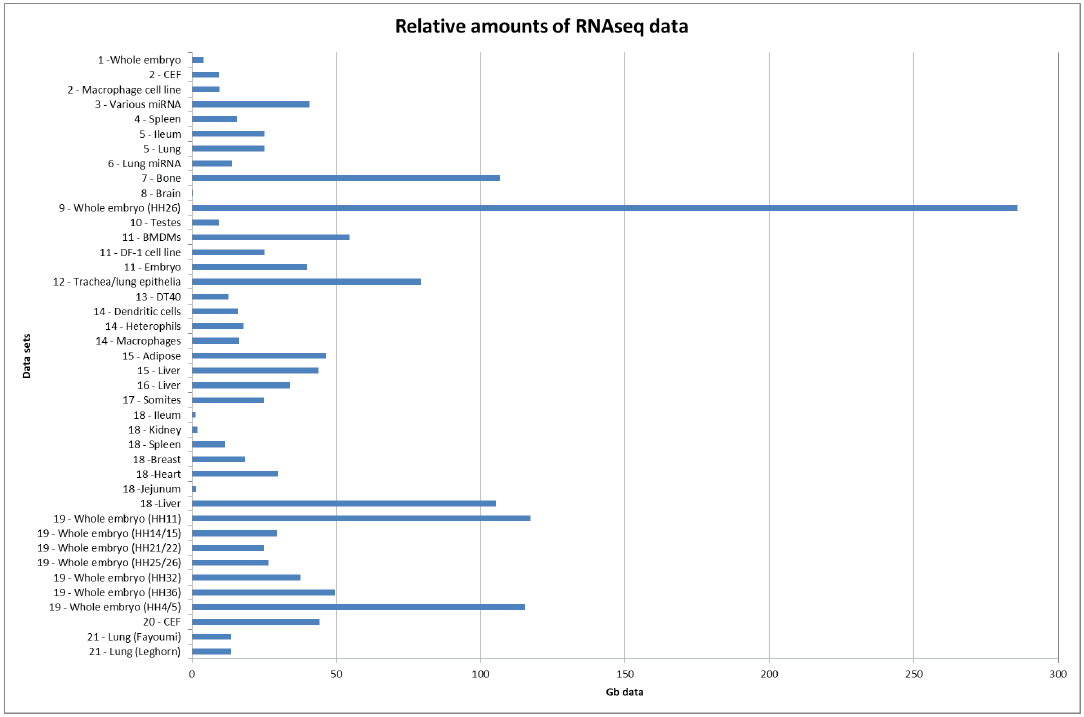
A comparison of the different relative amounts of RNAseq data from each tissue. Tissues from different data providers are shown separately as they have all been subject to different treatments/stimuli. Numbered data sets are as referred to in Table 2.

**Table 3:**
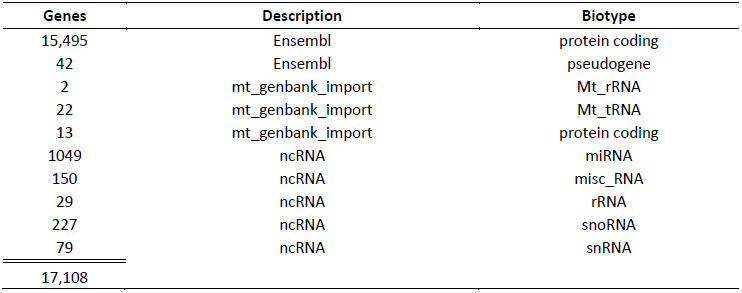
Ensembl 71 annotation statistics

**Table 4:** Comparison of Ensembl gene_builds:

|  | **Ensembl 70** | **Ensembl71** | **Ensembl 77** |
| --- | --- | --- | --- |
| **Assembly** | WashUC2, May 2006 | Galg4, Nov 2011 | Galg4, Nov 2011 |
| **Base pairs** | 1,050,947,331 | 1,072,544,086 | 1,072,544,763 |
| Coding genes | 16,736 | 15,508 | 15,508 |
| Short non-coding genes | 1,102 | 1,558 | 1,558 |
| Pseudogenes | 96 | 42 | 42 |
| Gene transcripts | 23,392 | 17,954 | 17,954 |

The availability of these data will allow for the further development of a chicken expression atlas by providing the ability to analyse transcript levels across tissues (http://geneatlas.arl.arizona.edu/). It will also enable development of exon capture technology for the chicken and has already proved of great use in helping annotate the other avian genomes which have now been sequenced. On-going collection of RNAseq data will remain a valuable resource as genomic analysis of avian species continues to expand.

## Methods

### Ensembl gene_build

The chicken gene_build from Ensembl release 71 was done using standard Ensembl annotation procedures and pipelines, mostly focussed on protein coding sequences. Briefly, vertebrate UniProtKB proteins were downloaded and aligned to the Galgal4 (GCA_000002315.2) assembly with Genewise (http://www.ebi.ac.uk/Tools/psa/genewise/) in order to annotate protein coding models. UniProt assigns protein existence (PE) levels to each of their protein sequences. The PE level indicates the type of evidence that supports the existence of a protein sequence, and can range from PE 1 (‘Experimental evidence at protein level’) to PE 5 (‘Protein uncertain’). Only PE 1 and PE 2 proteins from UniProtKB were used for the Genewise step. RNAseq models were annotated using the Ensembl RNAseq pipeline and models from both the Genewise and the RNAseq pipelines were used as input for the final protein-coding gene set. Chicken cDNAs and also RNAseq models were also used to add UTRs in the 5’ and 3’ regions. Some missing gene models were recovered by aligning chicken, zebra finch and turkey translations from Ensembl release 65 (December 2011) to the new chicken genome assembly.

### RNAseq Gene Models

Raw reads were aligned to the genome using BWA (Li & Durbin, 2009) to identify regions of the genome that are actively transcribed. The results from all tissues were used to create one set of alignment blocks roughly corresponding to exons. Read pairing information was used to group exons into approximate transcript structures called proto-transcripts. Next, partially mapped reads from both the merged (combined data from all tissue samples) and individual tissues were re-aligned to the proto-transcripts using Exonerate (Slater & Birney, 2005), to create a merged and tissue-specific sets of spliced alignments. For each gene, merged and tissue-specific transcript isoforms were computed from all observed exon-intron combinations, and only the best supported isoform was reported.

### Annotation of Non-Coding RNAs

The following non-coding RNA gene types were annotated - rRNA: ribosomal RNA; snRNA: small nuclear RNA; snoRNA: small nucleolar RNA; miRNA: microRNA precursors; misc_RNA: miscellaneous other RNA. Most ncRNA genes in Ensembl are annotated by first aligning genomic sequence against RFAM (Burge et al., 2013), using BLASTN (parameters W=12 B=10000 V=10000 -hspmax 0 -gspmax 0 -kap -cpus=1), to identify likely ncRNA loci. The BLAST (Altschul et al., 1990) hits are clustered, filtered for hits above 70% coverage, and used to seed an Infernal (Nawrocki & Eddy, 2013) search with the corresponding RFAM covariance model, to measure the probability that these targets can fold into the structures required. Infernal’s cmsearch is used to build ncRNA models. MiRNAs are predicted by BLASTN (default parameters) of genomic sequence slices against miRBase (Kozomara & Griffiths-Jones, 2014) sequences. The BLAST hits are clustered, filtered to select the alignment with the lowest p-value when more than one sequence aligns at the same genomic position, and the aligned genomic sequence is checked for possible secondary structure using RNAFold (Hofacker et al., 1994). If evidence is found that the genomic sequence could form a stable hairpin structure, the locus is used to create a miRNA gene model. Transfer RNAs (tRNAs) were annotated as part of the raw compute process using tRNAscan-SE with default parameters (Schattner et al., 2005). All results for tRNAscan-SE are available through Ensembl; the results are not included in the Ensembl gene set because they are not annotated using the standard evidence-based approach (ie. by aligning biological sequences to the genome) that is used to annotate other Ensembl gene models.

## Summary

The availability of this collection of chicken RNAseq data within the consortium has allowed:

- Annotation of 17,108 chicken genes, 15,495 of which are protein-coding (Ensembl 71)
- Identification of around 14,000 putative lncRNA genes (with >23,000 transcripts suggested)
- Annotation of miRNAs, snoRNAs, and other ncRNAs
- Future generation of an expression atlas which will allow comparisons of expression over many tissues
- An improved avian reference for comparative analyses with 48 other avian genomes (Zhang et al., 2014)

## Future directions

The next stage in progressing annotation of the avian genomes will concentrate on the analysis of data generated by PacBio sequencing, in conjunction with stranded RNAseq data from a wide variety of tissues. PacBio technology allows for very long read lengths, producing reads with average lengths of 4,200 to 8,500 bp, with the longest reads over 30,000 base pairs. This enables sequencing of full-length transcripts. Extremely high accuracy means that *de novo* assembly of genomes and detection of variants with greater than 99.999% accuracy is possible. Individual molecules can also be sequenced at 99% reliability. The high sensitivity of the method also means that minor variants can be detected even when they have a frequency of less than 0.1% [http://www.pacificbiosciences.com/products/smrt-technology/smrt-sequencing-advantage/]. We currently have brain transcriptomic PacBio data generated from a female Brown Leghorn J-line chicken (Blyth and Sang 1960). This will be analyzed alongside stranded RNAseq data that has been generated from 21 different tissues. The advantage of using strand-specific sequence information is that it provides an insight into antisense transcripts and their potential role in regulation and strand information of non-coding RNAs as well as aiding in accurately quantifying overlapping transcripts. It is particularly useful for finding unannotated genes and ncRNAs. This strategy should allow us to obtain full-length transcript sequences, identify novel transcripts and low-level transcripts, map transcription start and stop sites and confirm further ncRNAs.

## Get involved

If you’re interested in helping further the annotation of the avian genomes, and you can provide avian RNAseq data or can help with the analysis of such data, then please contact Jacqueline Smith or Dave Burt.

## Avian RNAseq Consortium Members

Jacqueline Smith, Ian Dunn, Valerie Garceau, David Hume, Pete Kaiser, Richard Kuo, Heather McCormack, Dave Burt (Roslin Institute); Amanda Cooksey, Fiona McCarthy, Parker B. Antin, Shane Burgess (University of Arizona); Andrea Münsterberg, Helio Pais (University of East Anglia); Andrew Oler (NIH National Institute of Allergy and Infectious Diseases); Steve Searle (Wellcome Trust Sanger Institute); Paul Flicek, Bronwen L. Aken, Rishi Nag (European Molecular Biology Laboratory, European Bioinformatics Institute and Wellcome Trust Sanger Institute); Carl Schmidt (University of Delaware); Christophe Klopp (INRA Toulouse); Pablo Prieta Barja, Ionas Erb, Darek Kedra, Cedric Notredame (CRG, Barcelona); David Froman (Oregon State University); Dirk-Jan de Koning (Swedish University of Agricultural Sciences, Uppsala); Douglas Rhoads (University of Arkansas); Igor Ulitsky (Weizmann Institute of Science, Rehovot); Julien Häsler, Michael Neuberger (*in memoriam*) (MRC, Cambridge); Laure Frésard, Frédérique Pitel (INRA, Auzville); Mario Fasold, Peter Stadler (University of Leipzig); Matt Schwartz (Harvard Medical School); Michael Skinner (Imperial College London); Olivier Hanotte (University of Nottingham); Perry Blackshear (NIEHS, North Carolina); Sandrine Lagarrigue, Pierre-François Roux (INRA Agrocampus Ouest); Thomas Derrien (University of Rennes); Sheila Ommeh (Jomo-Kenyatta University of Agriculture and Technology, Kenya); Stefan Muljo (NIH NIAID, Bethesda); Steve Kemp, Harry Noyes (University of Liverpool); Susan Lamont (Iowa State University); Ying Wang, Huaijun Zhou (UC Davis).

## Availability of RNASeq data

Data have been submitted to the public databases under the following accession numbers: Antin/Burgess/McCarthy/Schmidt data: BioProject ID: PRJNA204941 (Sequence Read Archive); Blackshear data: PRJEB1406 (European Nucleotide Archive); Burt/Smith data: E-MTAB-2908, E-MTAB-2909, E-MTAB-2910 (Array Express); De Koning/Dunn/McCormack data: E-MTAB-2737 (Array Express); Frésard/Pitel data: SRP033603 (Sequence Read Archive); Froman/Rhoads data: BioProject ID: PRJNA247673 (Sequence Read Archive); Garceau/Hume data: E-MTAB-3048 (Array Express); Hanotte/Kemp/Noyes/Ommeh data: E-MTAB-3068 (Array Express); Häsler/Oler/Muljo/Neuberger data: GSE58766 (NCBI GEO); Kaiser data: E-MTAB-2996 (Array Express); Lagarrigue/Roux data: SRP042257 (Sequence Read Archive); Lamont data: GSE51035 (NCBI GEO); Munsterberg/Pais data: GSE58766 (NCBI GEO); Schwartz/Ulitsky data: SRP041863 (Sequence Read Archive); Skinner data: PRJEB7620 (European Nucleotide Archive); Wang/Zhou data: GSM1385570, GSM1385571, GSM1385572, GSM1385573 (NCBI GEO).

